# Fecal Virome of *Coendou spinosus* (Paraguaian Hairy Dwarf Porcupine)

**DOI:** 10.1101/2023.09.24.559146

**Authors:** Thamiris dos Santos Miranda, Isabelle Gomes de Matos, Mirela D’arc, Filipe Romero Rebello Moreira, Francine Bittencourt Schiffler, Matheus Augusto Calvano Cosentino, Amanda Coimbra, Thiago Henrique de Oliveira, Ricardo Mouta, Gabriel Medeiros, Déa Luiza Girardi, Monique Lima, Victor Wanderkoke, Talitha Mayumi Francisco, Flávio Landim Soffiati, Suelen Sanches Ferreira, Carlos Ramon Ruiz-Miranda, Marcelo Alves Soares, André Felipe Andrade dos Santos

**Author notes:** These authors contributed equally to this work (Thamiris dos Santos Miranda, Isabelle Gomes de Matos).

## Abstract

The porcupine (*Coendou spinosus*) is a rodent species widely distributed in the Brazilian Atlantic forest. Like other rodent species, porcupines can be potential reservoirs of zoonotic agents. However, little is known about the viral diversity in these animals. Therefore, the objective of the present study was to evaluate, through massive sequencing, the virome of the feces of seven adult healthy free-living porcupines living in forest fragments from Silva Jardim, State of Rio de Janeiro, Brazil. A total of 41 viral families were classified, of which only seven were validated by both Kraken2 and Diamond taxonomic analysis tools, including bacteriophages (*Siphoviridae, Myoviridae, Podoviridae*), vertebrate viruses, such as *Papillomaviridae*, and unclassified RNA viruses. In addition, we also observed the presence of pathogenic bacteria and fungi already described in porcupines. The present study describes for the first time the microbiome in fecal samples from Brazilian porcupines, contributing to the global metagenomic characterization.

## MAIN TEXT

Porcupines (*Coendou spinosus*) (F. Cuvier, 1823) belong to the Erethizontidae family and are nocturnal arboreal Neotropical rodents, characterized by hair that is modified into sharp spines and prehensile tails (Menezes et al., 2020). Brazil has the greatest diversity of erectizontids spread in a wide range of habitats, including the Cerrado (Neotropical savannah), Pantanal and the Atlantic Forest (Voss et al., 2013). Some studies have already shown that rodents can be reservoirs of new viruses, which have been previously associated with emergence in human populations (Han et al., 2015; Gravinatti et al., 2020; Niendorf et al., 2021). Given the rise in frequency of epidemics caused by viruses of zoonotic origin, increasing research and surveillance efforts have been developed to characterize the viromes of wild fauna (A Duarte et al., 2019; Negrey et al., 2020). The aim of this study was to assess for the first time the virome of wild *Coendou spinosus* individuals sampled in Rio de Janeiro state, Brazil, through high-throughput sequencing (HTS).

The animals were captured between the months of May and July of 2019 in the municipality of Silva Jardim, state of Rio de Janeiro, Brazil (**Table 1**). The Igarapé farm belongs to the ecological park within the *Associação Mico Leão Dourado* (AMLD) and the other two farms belongs of the *Área de Proteção Ambiental da Bacia do Rio São João/Mico-Leão-Dourado* (APA). The porcupines were captured and samples manipulated under the approval and legal consent of the Brazilian Federal Authority for activities for scientific purposes under the numbers 67274-8 and 64635-5. Fecal samples were collected from seven adult healthy porcupines in the localities of Iguapê (CSP01, CSP03 and CSP09), Igarapé (CSP10, CSP11 and CSP13) and Flandria (CSP12), following national guidelines and provisions of CONCEA (Conselho Nacional de Controle de Experimentação Animal, Brazil). Samples were transferred to 15 mL falcon tubes with RNAlater™ (Thermo Fisher Scientific, Walham, USA) kept at room temperature and sent to the *Laboratório de Diversidade e Doenças Virais* (LDDV) of the *Universidade Federal do Rio de Janeiro* (UFRJ), to be stored at −80 ºC.

**Table 1:** General sampling information.

| ID | Sex | Location | GPS* |
| --- | --- | --- | --- |
| CSP01 | M | Fazenda Iguapê, Silva Jardim, RJ | -22.505761, -42.323408 |
| CSP03 | F | Fazenda Iguapê, Silva Jardim, RJ | -22.505761, -42.323408 |
| CSP09 | F | Fazenda Iguapê, Silva Jardim, RJ | -22.505761, -42.323408 |
| CSP10 | M | Fazenda Igarapé, Silva Jardim, RJ | -22.50713, -42.30954 |
| CSP11 | F | Fazenda Igarapé, Silva Jardim, RJ | -22.50713, -42.30954 |
| CSP12 | F | Fazenda Flandria, Silva Jardim, RJ | -22.50950, -42.31868 |
| CSP13 | F | Fazenda Igarapé, Silva Jardim, RJ | -22.50713, -42.30954 |

\*Global Positioning System in Decimal Degrees format

The next steps were viral metagenomics library construction, sequencing and computational analysis. Briefly, samples went through a viral enrichment step, followed by an indexed paired-end library preparation with the Nextera XT kit, according to the manufacturer’s instructions with some modifications described previously (Cosentino et al., 2022). Sequencing was performed on the MiSeq instrument (Illumina) with 300 cycles per sequencing pair-ended (2 × 151 bp) from the Department of Genetics at UFRJ. An *in house* custom bioinformatics pipeline was used for data quality control with Fastp v.0.20.1 (Chen et al., 2018), *de novo* assembly using SPAdes v3.15.3 (Nurk et al., 2017), taxonomic assignments were performed by Kraken2 (Wood & Salzberg, 2014) and Diamond v.2.0.14 (Buchfink et al., 2015) and interactive visualization with Krona v.2.7.1 (Ondov et al., 2011), also described previously in Cosentino *et al*., (2022). The library generated a total of 2,109,411 raw reads, of which 949,294 reads (45%) passed the criteria for quality trimming and filtering. Through Kraken2, a total of 350,204 reads (37%) were taxonomically classified. Of this total, the highest proportion of reads were Bacteria (192,280 reads; 54.8%), followed by Eukaria (156,712 reads; 44.7%) and a small number of viral sequences (344 reads; 0.1%) (**Supplementary Table S1**). After cutting off 1% contamination between libraries from the same run, 23 viral families were classified (**Supplementary Table S2**).

A total of 147,187 reads (15.5%) were taxonomically assigned with the Diamond tool (NCBI nr database). Most classified reads were assigned to bacteria (135,586 reads; 92.2%), followed by eukaryotes (10,723 reads; 7.3%) and a small percentage of viruses (475 reads; 0.3%) **(Figure 1a**; **Supplementary Table S1**). When analyzing bacterial data, we observed the presence of potential bacterial pathogens already described in hedgehogs, such as *Anaplasma, Lepstopira, Mycobacterium, Staphylococcus, Streptococcus* and *Corynebacterium*, the last four of which can cause diseases in porcupines (Ruszkowski et al., 2021) (**Supplementary File S1**). We also observed few sequences taxonomically related to fungi such as *Candida*, which has already been demonstrated that porcupines can be a potential carrier or host of these fungi that are pathogenic in humans (Barlow et al., 2012), but further studies are needed to validate the presence of this fungus in these samples.

**Figure 1:**
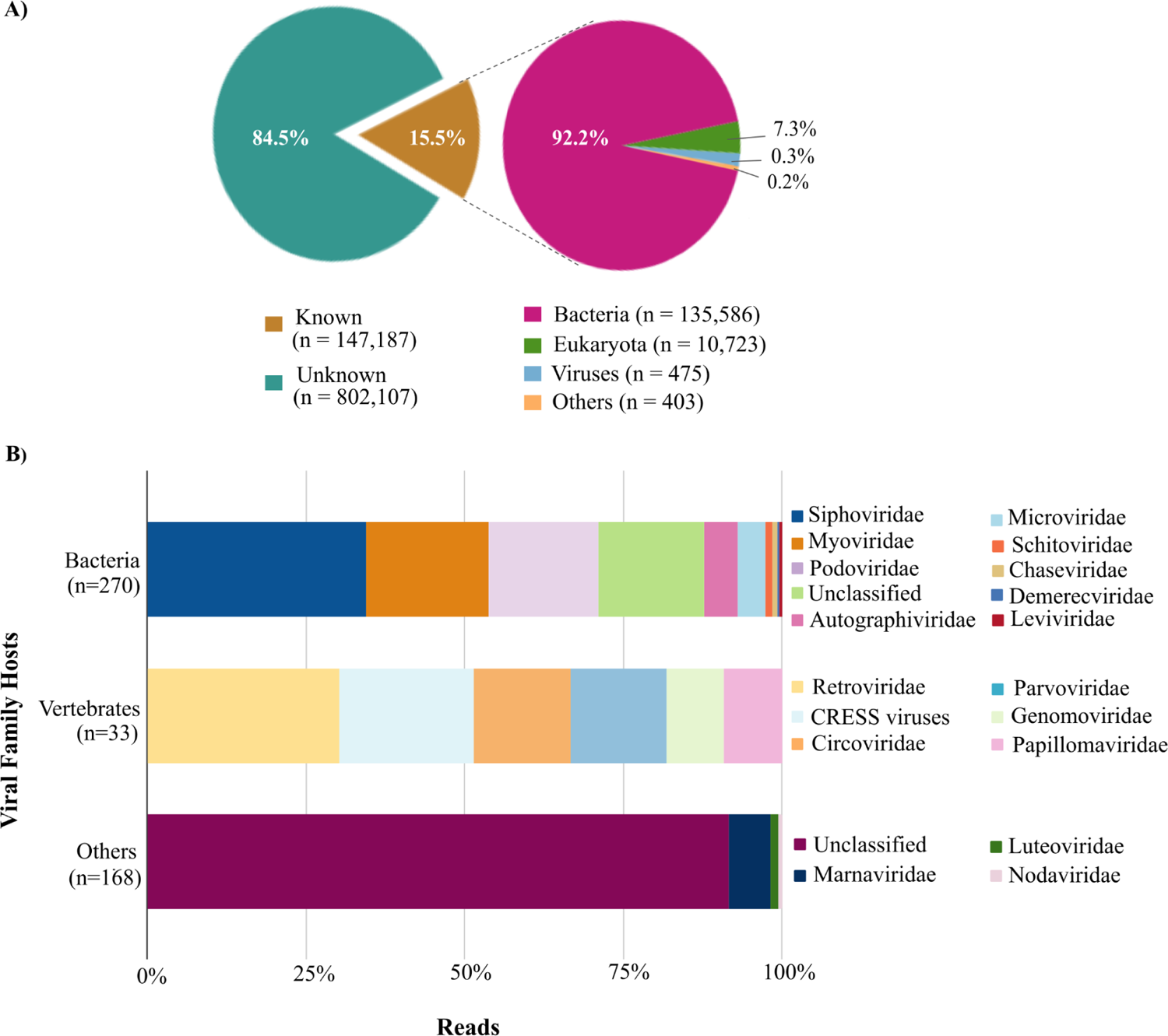
Taxonomic classification of sequences in porcupine samples by Diamond. (A) The percent of “known” and “unknown” virome reads when compared to the protein database (NCBI nr database) and classification of “known” sequences in Bacteria, Eukarya, Viruses and Others (including Archaea and unassigned sequences). (B) Taxonomic composition at the viral family level grouped by host type like Bacteria, Vertebrates and Others (including protists, plants, invertebrates and unclassified viruses). The different families were denoted with different colors.

Among the viral reads, a total of 18 viral families were confirmed, with bacteriophages groups being the most abundant, as *Siphoviridae* (93 reads; 19.74%), *Myoviridae* (52 reads; 11.04%), *Podoviridae* (47 reads; 9.97%) and *Autographiviridae* (14 reads; 2.97%) (**Figure 1b; Supplementary Table S2)**. Vertebrate viruses such as *Retroviridae* (10 reads; 2.12%), unidentified CRESS virus (7 reads; 1.48%), *Circoviridae* (5 reads; 1.06%), *Parvoviridae* (5 reads; 1.06%), *Genomoviridae* (3 reads; 0.68%) and *Papillomaviridae* (3 reads; 0.68%), were also classified. Other viral families found in protozoans and plants have also been identified, *Marnaviridae* (11 reads; 2.33%) and *Luteoviridae* (2 reads; 0.42%) respectively. Through *de novo* assembly we obtained a total of 4,107 contigs (ranging from 125 - 2,067 nt). However, only three contigs were classified for the viral families *Siphoviridae, Podoviridae* and *Demerecviridae* by Kraken2 and two contigs for the *Retroviridae* family and unclassified DNA virus by Diamond.

In all, about 62.8% DNA viruses and 37.2% RNA viruses were sequenced when considering the two taxonomic analyses. Among the viral families identified by Diamond, only seven (38.8%) were concordant with Kraken2 (standard database) analysis. Most were bacteriophages such as *Siphoviridae, Myoviridae, Podoviridae, Autographiviridae, Schitoviridae* and *Demerecviridae*, commonly found in the feces of animals and humans (Chong et al., 2019; Liang & Bushman, 2021; Ning et al., 2021) (**Supplementary Table S2**). The only vertebrate viral family validated by both tools was *Papillomaviridae*, often associated with various benign and malignant epithelial lesions in their natural host, including porcupines (Elfadl et al., 2017; Shimazaki et al., 2023). In addition to the seven viral families, a large proportion of unclassified ribovirus were classified by Kraken2 (22 reads; 6.4%) and Diamond (136 reads; 28.9%) (**Supplementary Table S2**). These viral reads were investigated in greater detail by reference mapping to the most closely related viral genome (NCBI accession number: NC_032884) with Geneious software v2021.2.2 using default parameters (Kearse et al., 2012). Through this method, we obtained seven contigs (138-1,651 pb) with a coverage in the reference genome of 87.8%. BLASTn of the consensus sequence showed that it had a high nt identity (74.2%) to *Hubei tombus-like virus 8 strain* (accession: KX883226.1). Using the fgenesVO (www.softberry.com) to find ORFs in viral genomes, an ORF of 235 aa was found and shown by BLASTp to have 100% of coverage and 73% of identity for the hypothetical protein 3 from *Hubei tombus-like virus 8* (YP_009336791.1), described in insects. Although Hubei tombus-like virus has been commonly classified as a plant virus, this definition has changed recently when unassigned *Hubei tombus-like viruses* (8 and 20) were discovered in invertebrate animals (Reuter et al., 2019). We believe this is one more instance where analysis of the fecal virome led to the identification of a previously unknown dietary-related virus (Shi et al. 2016; Reuter et al. 2019; Miranda et al. 2023).

Viral metagenomics has recently been employed in numerous studies, providing insight into the composition of animal viromes and helping to identify novel viruses, including agents with zoonotic transmission potential (Chand, 2022; He et al., 2022). The wild porcupine can be an important vector of viral, bacterial and fungi agents, such as flaviviruses and poxviruses (Thoisy et al., 2003, 2004; Hora et al., 2021; Guerra et al., 2022;). None of these viral pathogens were detected in this study, consistent with the healthy status of all animals at the time of sample collection. In conclusion, this study assessed for the first time the virome of fecal samples from Brazilian porcupines, providing evidence for the circulation of multiple viral families in animals of the Erethizontidae family. Furthermore it was possible to discover the presence of an unassigned group of RNA viruses. This study exemplifies how virome analysis from non-invasive samples can be used to monitor wildlife health.

## DATA AVAILABILITY

Raw sequencing reads are available under BioProject accession number PRJNA972708 and NCBI SRA accession number SRR24628734. Assembled contigs, taxonomic assignment files and Krona plots are available on the project GitHub page (link: https://github.com/lddv-ufrj/Porcupine_Virome).

## ACKNOWLEDGMENTS

Field work was supported by resources from PETROBRAS (process 2017/00606-7) to Carlos R Ruiz-Miranda for the project Evaluation of the effect of pipeline tracks on landscape connectivity for Mammals and analysis of the effectiveness of fauna crossing structures. This work is also supported by the National Council for Scientific and Technological Development/CNPq (grant number 313005/2020-6 for AFS) and Fundação de Amparo à Pesquisa do Estado do Rio de Janeiro/FAPERJ (grant number E26/201.193/2022, E26/211.355/2021 and E26/211.040/2019 for AFS). We are grateful for the use of Molecular Biology equipment at the Common Use and Technical Support Laboratory of the Federal University of Rio de Janeiro (Rio de Janeiro, Brazil) and all collaborators for their help in sample collection, especially Caique Ferreira Amaral Soares.

## CONFLICTS OF INTEREST

The authors declare no conflict of interest.

## Notes

https://github.com/lddv-ufrj/Porcupine_Virome

### Competing Interest Statement

The authors have declared no competing interest.

### Summary of Updates

Author affiliations updated

