## Supplementary Table S1 for "Fecal Virome of *Coendou spinosus* (Paraguaian Hairy Dwarf Porcupine)"

**Supplementary Table S1:** Virome overview.

| | | | |
|---|---|---|---|
| **Raw reads** | | 2,109,411 | |
| **Filtered reads** | | 949,294 | |
| **G+C content (%)** | | 48% | |
| | | **Kraken2** | **Diamond** |
| **No hits** | **Reads (%)** | 598,442 (63%) | 802,107 (84.5%) |
| | **Contigs (%)<br>(min - max)** | 2,266 (55%)<br>(125 - 1,703 nt) | 1,049 (25.5%)<br>(125 - 1,625 nt) |
| **Mapped reads** | | 350,852 (37%) | 147,187 (15.5%) |
| **Mapped contigs<br>(min - max)** | | 1,841<br>(129 - 2,067 nt) | 3,058<br>(125 -2,067 nt) |
| **Bacteria** | **Reads (%)** | 192,280 (54.8%) | 135,586 (92.2%) |
| | **Contigs (%)<br>(min - max)** | 1,658 (90%)<br>(375 - 2,067 nt) | 2,715 (88.8%)<br>(125 - 2,067 nt) |
| **Eukaryota** | **Reads (%)** | 156,712 (44.7%) | 10,723 (7.3%) |
| | **Contigs (%)<br>(min - max)** | 167 (9.1%)<br>(129 - 1,538 nt) | 338 (11.04%)<br>(375 - 1,703 nt) |
| **Viral** | **Reads (%)** | 344 (0.1%) | 475 (0.3%) |
| | **Contigs (%)<br>(min - max)** | 3 (0.2%)<br>(436 - 545 nt) | 2 (0.06%)<br>(407 - 440 nt) |
| **Others/<br>Unassigned** | **Reads (%)** | 1,516 (0.4%) | 403 (0.2%) |
| | **Contigs (%)<br>(min - max)** | 13 (0.7%)<br>(388 - 831 nt) | 3 (0.1%)<br>(387 - 673 nt) |
