## Supplementary Table S2 for "Fecal Virome of *Coendou spinosus* (Paraguaian Hairy Dwarf Porcupine)"

**Supplementary Table S2:** Viral diversity founded in Kraken2 and Diamond analysis.

| VIRUS FAMILY | KNOWN HOSTS | GENOME TYPE | KRAKEN2 | | DIAMOND | |
|---|---|---|---|---|---|---|
| | | | **N° Viral Reads** | **N° Viral Contigs** | **N° Viral Reads** | **N° Viral Contigs** |
| ***Myoviridae**** | Bacteria | **dsDNA-L** | 86 | 0 | 52 | 0 |
| ***Siphoviridae**** | Bacteria | **dsDNA-L** | 74 | 1 | 93 | 0 |
| ***Podoviridae**** | Bacteria | **dsDNA-L** | 30 | 1 | 47 | 0 |
| ***Phycodnaviridae*** | Algae | **dsDNA-L** | 20 | 0 | 0 | 0 |
| ***Baculoviridae*** | Invertebrates | **dsDNA-C** | 16 | 0 | 0 | 0 |
| ***Herelleviridae*** | Bacteria | **dsDNA-L** | 14 | 0 | 0 | 0 |
| ***Autographiviridae**** | Bacteria | **dsDNA-L** | 8 | 0 | 14 | 0 |
| ***Iridoviridae*** | Vertebrates, Invertebrates | **dsDNA-L** | 8 | 0 | 0 | 0 |
| ***Demerecviridae**** | Bacteria | **dsDNA-L** | 6 | 1 | 1 | 0 |
| ***Papillomaviridae**** | Vertebrates | **dsDNA-C** | 6 | 0 | 3 | 0 |
| ***Poxviridae*** | Vertebrates | **dsDNA-L** | 6 | 0 | 0 | 0 |
| ***Reoviridae*** | Vertebrate, Invertebrate, Plant, Fungi | **dsRNA-L** | 6 | 0 | 0 | 0 |
| ***Nimaviridae*** | Invertebrates | **dsDNA-C** | 4 | 0 | 0 | 0 |
| ***Drexlerviridae*** | Bacteria | **dsDNA-L** | 4 | 0 | 0 | 0 |
| ***Schitoviridae**** | Bacteria | **dsDNA-L** | 2 | 0 | 3 | 0 |
| ***Coronaviridae*** | Vertebrates | **ssRNA-L** | 2 | 0 | 0 | 0 |
| ***Arenaviridae*** | Vertebrates | **ssRNA-L** | 2 | 0 | 0 | 0 |
| ***Arteriviridae*** | Vertebrates | **ssRNA-L** | 2 | 0 | 0 | 0 |
| ***Potyviridae*** | Plants | **ssRNA-L** | 2 | 0 | 0 | 0 |
| ***Bromoviridae*** | Plants | **ssRNA-L** | 2 | 0 | 0 | 0 |
| ***Lavidaviridae*** | Protists | **dsDNA-C** | 2 | 0 | 0 | 0 |
| ***Mimiviridae*** | Protists | **dsDNA-L** | 2 | 0 | 0 | 0 |
| ***Pospiviroidae*** | Plants | **ssRNA-C** | 2 | 0 | 0 | 0 |
| ***Microviridae*** | Bacteria | **ssDNA-C** | 0 | 0 | 12 | 0 |
| ***Chaseviridae*** | Bacteria | **dsDNA-L** | 0 | 0 | 2 | 0 |
| ***Leviviridae*** | Bacteria | **ssRNA-L** | 0 | 0 | 1 | 0 |
| ***Retroviridae*** | Vertebrates | **ssRNA-L** | 0 | 0 | 10 | 1 |
| **CRESS viruses** | Vertebrates | **ssDNA-C** | 0 | 0 | 7 | 0 |

| | | | | | | |
|---|---|---|---|---|---|---|
| ***Circoviridae*** | Vertebrates | **ssDNA-C** | 0 | 0 | 5 | 0 |
| ***Parvoviridae*** | Vertebrates | **ssDNA-L** | 0 | 0 | 5 | 0 |
| ***Genomoviridae*** | Vertebrates, Invertebrates | **ssDNA-C** | 0 | 0 | 3 | 0 |
| ***Marnaviridae*** | Protists | **ssRNA-L** | 0 | 0 | 11 | 0 |
| ***Luteoviridae*** | Plants | **ssRNA-L** | 0 | 0 | 2 | 0 |
| ***Nodaviridae*** | Vertebrates, Invertebrates | **ssRNA-L** | 0 | 0 | 1 | 0 |
| ***Ligamenvirales*** | Bacteria, Archea | **dsDNA-L** | 4 | 0 | 0 | 0 |
| ***Caudovirales*** | Bacteria, Archea | **dsDNA-L** | 4 | 0 | 1 | 0 |
| ***Megaviricetes*** | Protists, Vertebrates, Invertebrates | **dsRNA-L** | 0 | 0 | 5 | 0 |
| ***Unclassified Picornavirales*** | Vertebrates | **ssRNA-L** | 0 | 0 | 1 | 0 |
| ***Unclassified bacterial viruses*** | Bacteria | **ssRNA-L** | 0 | 0 | 6 | 0 |
| ***Unclassified Cressdnaviricota*** | Bacteria, Animals, Plants | **ssDNA-C** | 0 | 0 | 1 | 0 |
| ***Unclassified Riboviria*** | Plants, Invertebrates | **RNA** | 22 | 0 | 136 | 0 |
| ***Unclassified viruses*** | Bacteria | **DNA** | 8 | 0 | 38 | 1 |
| ***Environmental samples*** | Bacteria, Invertebrates, Vertebrates, Plants, Protists | **RNA** | 0 | 0 | 11 | 0 |
| **Total** | | | 344 | 3 | 471 | 2 |

* Viral families concordant with Kraken2 and Diamond.
